# Multifunctional bending magnet beamline with a capillary optic for X-ray fluorescence studies of metals in tissue sections

**DOI:** 10.1101/2025.01.18.633695

**Authors:** Benjamin Roter, Andrew M. Crawford, Qiaoling Jin, Arthur T. Glowacki, Barry Lai, Fabricio S. Marin, Evan Maxey, Xianbo Shi, Valeria C. Culotta, Asia S. Wildeman, Naisargi K. Patel, Thomas V. O’halloran, Chris Jacobsen

**Author notes:** These authors contributed equally to this work (Benjamin Roter, Andrew M. Crawford).

## Abstract

Scanning fluorescence X-ray microscopy lets one non-destructively and quantitatively map the distribution of most biologically-important metals in cells and tissues. For studies on large-scale tissues and organs, a spatial resolution of several micrometers is often sufficient; in this case, bending magnets at synchrotron light sources provide abundant X-ray flux. We describe here the use of bending magnet beamline 8-BM-B at the Advanced Photon Source (APS) with two distinct microscopy stations: a pre-existing one with Kirkpatrick-Baez (KB) mirror optics for slightly higher throughput and the ability to accommodate samples tens of centimeters across, and a new prototype station with an axially-symmetric, single-bounce, capillary optic with slightly less flux, but slightly higher fluence (which affects achievable resolution at low metal concentration) and higher spatial resolution. The KB station provides *δ*_res_ = 10.5 µm spatial resolution at a per-pixel exposure time of *t*_dwell_ = 100 ms and a fluence per time of 5.8× 10^7^ photons /(µm^2^ ·s), while the prototype capillary station provides *δ*_res_ = 6.3 µm at *t*_dwell_ = 50 ms and a fluence per time of 6.1× 10^7^ photons (µm^2^ ·s). We used image power spectral density to estimate the achieved spatial resolution *δ*_res_ from individually acquired images, with *δ*_res_ depending-on the optic, the fluorescence signal strength of the sample being imaged, and the method used to process raw fluorescence spectral data.

## 1. Introduction

Quantitative methods for mapping elemental distributions are essential for addressing a wide range of problems in the life sciences [1, 2]. To understand the function of metals in thin sections from tissues and organs, quantitative imaging at a spatial resolution of several micrometers can provide information on cell-to-cell metal variations, while a field of view of several millimeters allows one to image large, representative regions of organs from small animals. For elements with atomic numbers above about *Z* = 14, scanning fluorescence X-ray microscopy (SFXM) offers a very useful combination of high sensitivity with relatively low beam damage [3–5]. In this approach, a small specimen is raster-scanned through a X-ray beam spot, and an energy-dispersive detector is used to record the emission at characteristic X-ray fluorescence lines [4, 6, 7]. The recorded signal includes background from elastic and Compton X-ray scattering, as well as other factors such as incomplete charge collection in energy-dispersive detectors [8]. A variety of analysis software packages can be used to separate signal from background, delivering quantitative measures of elemental concentration [9–14].

Obtaining high resolution in SFXM requires an excitation X-ray source providing significant flux within a small area with narrow solid angle; that is, a source with high spectral brightness [15]. Emission from such a source can then be demagnified by an optic to a small spot through which the sample is scanned. For nanoscale imaging within single cells, undulator sources in straight sections of low-emittance storage rings provide the highest possible brightness [16] outside of free-electron lasers. When evaluating overall elemental distributions in biological tissues or organs, micrometer-scale imaging can provide megapixel images of samples that are millimeters in size. In this case, the somewhat larger source size and divergence of bending magnet (dipole) sources at synchrotron light sources can provide quite satisfactory performance, often with easier access for imaging many samples.

We describe here the use of bending magnet beamline 8-BM-B at the Advanced Photon Source at Argonne National Laboratory. This beamline has a pre-existing SFXM station that uses a Kirkpatrick-Baez (KB) mirror [17] originally designed for use at a separate synchrotron beamline. As part of a project for Quantitative Elemental Mapping for the Life Sciences (QE-MAP), which is an NIH-supported Biomedical Technology Research Resource based at Michigan State University, we have added a second prototype SFXM station just downstream that is equipped with both a capillary focusing optic for higher spatial resolution and a fluorescence detector with greater signal collection capability. These two stations will be described in more detail in Sec. 1.2.

### 1.1 Example: Metals in a mouse kidney

As an example of SFXM capabilities provided at beamline 8-BM-B, Fig. 1 shows the distribution of three important metals in a mouse kidney partial section. This particular mouse was infected with *Candida albicans* clinical isolate SC5314 via tail vein injections as part of a larger study aimed at understanding how mammalian hosts attempt to starve invading pathogens of essential nutrient metals such as Fe, Cu, Zn, and Mn [18]. At 72 hours post-infection, the kidney was extracted, embedded in an optimum cutting temperature (OCT) compound (Tissue-Tek), and frozen in an isopentane bath chilled with liquid N_2_-cooled isopentane. The frozen block was cut into sections of 10 µm thickness while using a cryostat (CM3050S, Leica) that maintained them at −15-18°C during the process. The sections were transferred to Si_3_N_4_ chips (NX5200, Norcada). These chips were previously affixed to glass slides by Kapton tape. We then stored the sections in airtight slide boxes in a −80°C freezer until retrieval for SFXM analysis. Upon thawing the original airtight slide boxes at room temperature, we removed the chips containing the kidney sections from the glass slides and placed them onto a sample mount specifically designed for 8-BM-B (at which point most of the water in the sections evaporated). We scanned those sections at 8-BM-B, using an AXO 10X thin film standard (Applied X-ray Optics, GmbH) scanned separately to calibrate the metal concentrations. We obtained two-dimensional elemental maps of each section by performing fluorescence spectrum fitting with MAPS [10] and M-BLANK [14] software as will be described in Sec. 1.5. In Fig. 1(a), elemental maps of iron, calcium, and zinc are shown as obtained using the KB mirror station with a scan step size of Δ_*x*_ = Δ_*y*_ = 25 µm and a per-pixel exposure time *t*_dwell_ = 50 ms. Also shown in Fig. 1(a) is a three-color composite map of these three elements. The ring-like features in the Ca distribution indicate possible localized infection sites. Figs. 1(b) and (c) show finer fields of view of localized Ca hotspots at different length scales using (b) the KB mirror with Δ_*x*_ = Δ_*y*_ = 5 µm and *t*_dwell_ = 100 ms, and (c) the capillary optics with Δ_*x*_ = 1 µm, Δ_*y*_ = 2 µm in the original sample orientation, and *t*_dwell_ = 50 ms. As can be seen, the capillary optic station yields a higher resolution image.

**Fig. 1.**
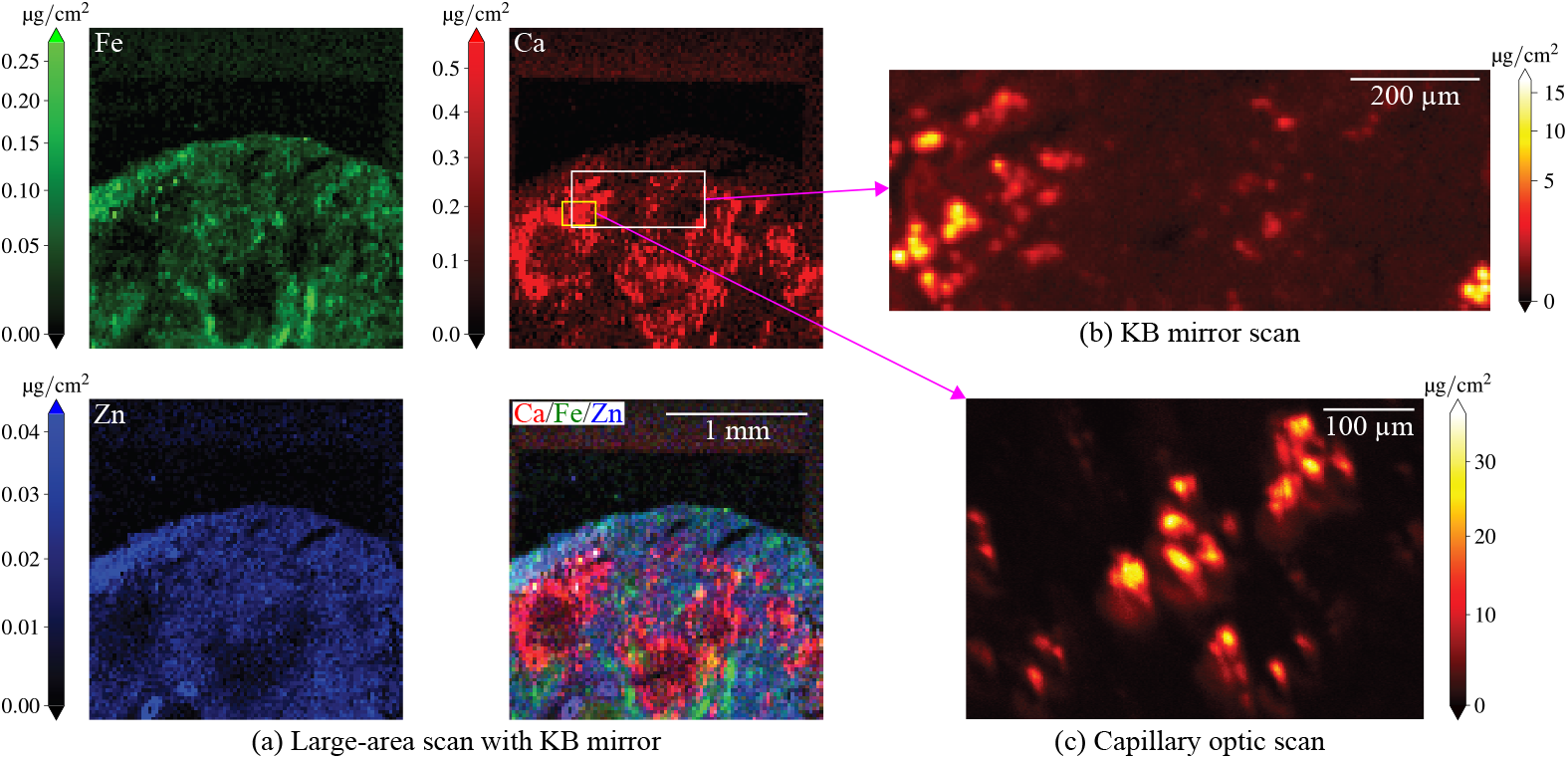
Scanning fluorescence X-ray microscopy (SFXM) images elemental distributions at different length scales. Shown here are trace essential metals inside a sample of a mouse kidney partial section infected with *Candida albicans*, with Ca being strongly elevated at sites of infection. We obtained the images in (a) using the KB mirror scanning station at 8-BM with a 25 µm step size at *t*_dwell_ = 50 ms. These metals are also displayed in a color composite map, allowing one to more easily see how the elements are differentially distributed in the specimen. The images in (b) and (c) correspond to regions we selected from (a) using the KB mirror and capillary optics, respectively. In (b), we used a Δ_*x*_ = Δ_*y*_ = 5 µm step size at *t*_dwell_ = 100 ms. In (c), we used Δ_*x*_ = 1 µm step size in *x* and Δ_*y*_ = 2 µm step size in *y* (in the original sample scan orientation) at *t*_dwell_ = 50 ms. The spatial resolution improved from 10.5 µm to 6.3 µm when transitioning from the KB optic to the capillary (as described in Sec. 2.2). We arranged the capillary image shown in (c) to match the images obtained with the KB mirror. The large-scanning area SFXM capabilities of 8-BM described in this work provide an important complement to nanoscale imaging capabilities from other beamlines at the Advanced Photon Source at Argonne National Laboratory.

### 1.2 Beamline 8-BM-B at the Advanced Photon Source

Beamline 8-BM-B at the Advanced Photon Source (APS) has had an evolving mission, with corresponding changes in its instrumentation. This bending magnet beamline was built in the mid-1990s as a general purpose beamline. Around 2010, it was repurposed for micrometer-scale X-ray microprobe studies. The present beamline optical layout is shown in Fig. 2. A double-multilayer monochromator (DMM) located 25.25 m from the source is used to deliver a spectral bandwidth of about Δ*λ*/ *λ* = 0.0109 at a photon energy of 10 keV, and a set of secondary source apertures are located 48.15 m from the primary source, just inside the experimental enclosure (the “hutch”). A vertically-deflecting, bendable toroidal mirror (originally designed for a different beamline) is located 30.9 m from the source; it delivers an intermediate focus spot located significantly upstream of the beam-defining aperture (BDA). Within the hutch, the secondary source apertures are located 0.2 m from the upstream end of a 2.4 m long optical table. At the table’s downstream end is an X-ray beam camera consisting of a 1 mm thick cadmium tungstate scintillator (10 mm side length), a 5× microscope objective, and a visible light camera. This camera is used for optic and specimen alignment. The BDA consists of a 250 µm diameter hole in a 0.25 mm thick platinum plate in front of a 1 mm diameter hole, which is in turn located inside a 12.7 mm thick aluminum plate. At the time of our measurements, the beam profile was roughly (1 mm) × (2 mm) at the input of the pinhole. There is the potential to enhance beamline performance by upgrading the toroidal mirror, and the source characteristics are being improved by the realization of the APS-Upgrade (APS-U) [19].

**Fig. 2.**
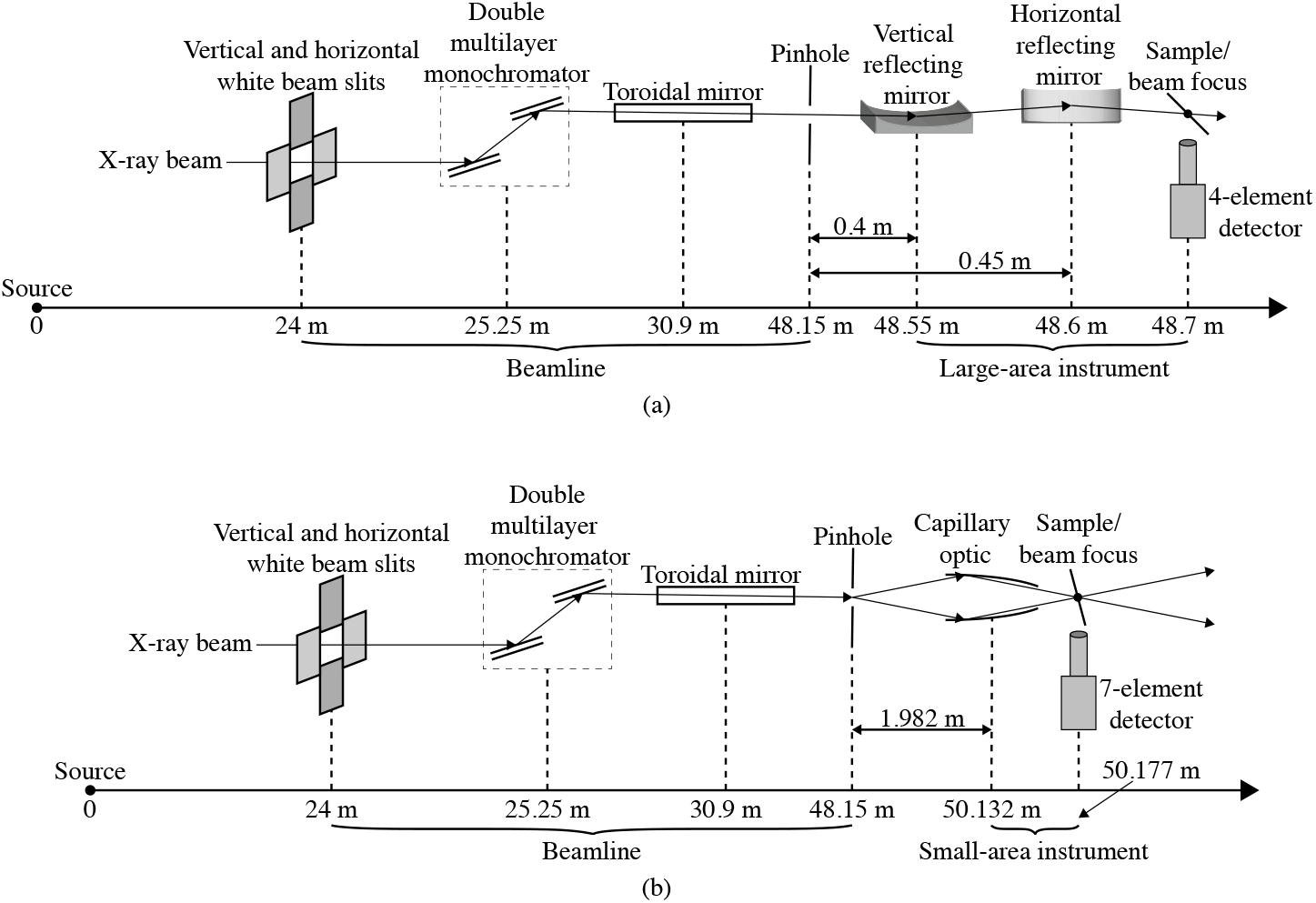
Layout of the 8-BM-B beamline when used with both a pre-existing KB mirror/large-area instrument setup (a) and a new, additional capillary/small-area instrument (b) located in the experimental enclosure. Both schematics show the location of the double multilayer monochromator (DMM), and the toroidal mirror intended to image the X-ray beam from the storage ring to the exit slit position. The beam size is then set using a pinhole as a beam-defining aperture (BDA). The KB mirror setup is for scanning sample fields up to several centimeters across at about *δ*_res_ = 10.5 µm spatial resolution, and the capillary optic setup is for *δ*_res_ = 6.3 µm spatial resolution over smaller fields of view (Sec. 2.1).

This beamline already features a large-scanning-area SFXM setup employing a Kirkpatrick-Baez (KB) focusing optic. The KB scanning setup has a large working distance of about 50 mm, and the sample is mounted at 45° relative to the incident beam normal to allow for clearance for very large specimens and good X-ray fluorescence signal collection efficiency. The system is equipped with a SII NanoTechnology Vortex ME-4 silicon drift detector, with XIA XMAP detector readout electronics. Because of the imperfect match between 8-BM-B beamline optics and the KB mirror, the achieved resolution is reduced, as will be seen in Sec. 2.1. Nevertheless, the KB scan system has proven to be quite successful in studies involving large-area samples [20–23]. As noted in Sec. 1, we have enhanced the beamline by adding a second SFXM setup just downstream of the KB setup. This second scanning station, which is presently in a prototype rather than final status, uses an axially-symmetric, single-bounce, capillary focusing optic designed to deliver a smaller focus with the optical layout of the 8-BM beamline. This capillary was pulled under heating conditions by Sigray, Inc. with the goal of imaging a source located 1.982 m upstream to a focus located 45 mm downstream of the capillary center while maintaining a working distance of 20 mm. We evaluated the performance of this optic using the 28-ID-B undulator beamline at the APS so as to provide a well-collimated X-ray source; with this source, the optic produced a focal spot with a full width at half maximum (FWHM) size of *δ*_res,*x*_ = 2.9 µm and *δ*_res,*y*_ = 3.0 µm at *E* = 13 keV (see Supplement 1 Sec. S1 for more details). This optic is mounted on a stage with two-axis translation and tilt for alignment to the incident beam.

Following the optic is a micrometer-precision scanning stage with a scanning direction oriented at 15° relative to the incident beam normal (to reduce horizontal beam spreading on the sample; see Fig. 3), as well as a new 7-element energy-dispersive detector (SII NanoTechnology Vortex ME-7 SDD) with electronics capable of handling higher count rates (Quantum Detectors Xspress3X). Because of the smaller working distance of this optic and the 15° scan direction, this setup is better suited to use with samples with no upstream clearance problems. Sections from biological tissues and organs usually satisfy these requirements.

**Fig. 3.**
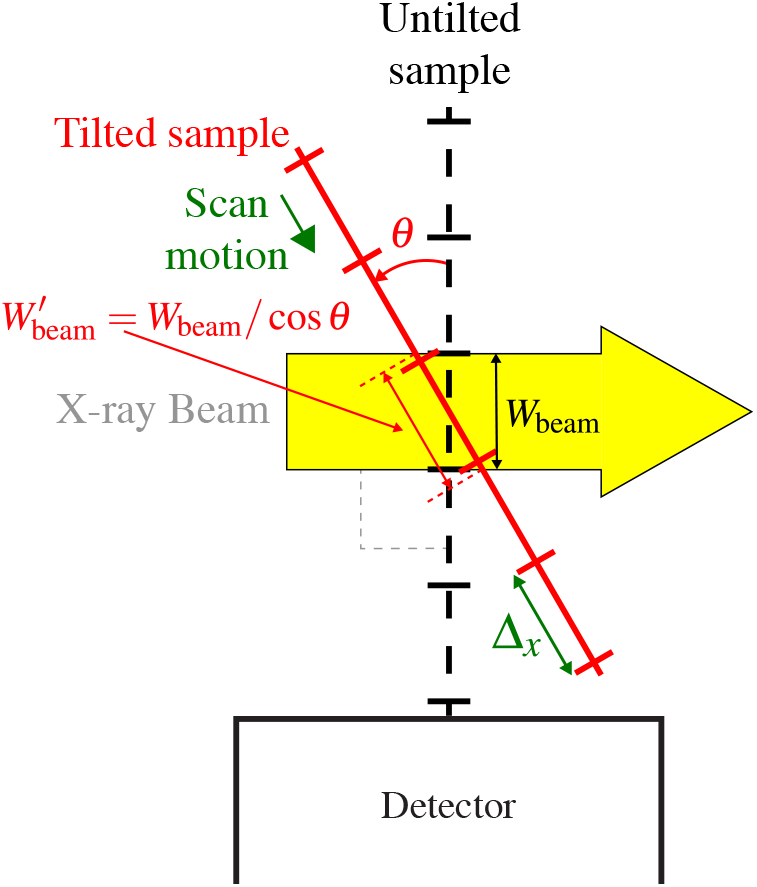
Diagram of how a planar sample and its scanning stage are typically mounted at an angle *θ* relative to normal incidence for scanning X-ray fluorescence microscopy. This means that the beam width *W*_beam_ as seen by a sample at normal incidence is broadened to 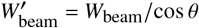.

With these two SFXM stations, the beamline 8-BM now has two scanning options. One is for larger samples at modest resolution, but with somewhat higher flux, using the KB optic with larger working distance. The other is for smaller samples using the capillary optic with reduced working distance, but higher spatial resolution and slightly increased fluence ℱ despite a slight decrease in flux Φ (see Sec. 3). The capillary station is presently in prototype status, with further improvements in discussion. Yet already biomedical and other users are able to choose which of these two scanning stations are best suited to their research interests without significant downtime for reconfiguring the SFXM system. This is quite useful at a facility where nanofocusing beamlines (which are often oversubscribed) prioritize users needing to collect data on biological samples at submicron resolution.

The achieved spatial resolution in SFXM depends not only on the properties of the optic, but also on the signal and background level for detecting particular chemical elements. While this has been illustrated previously for SFXM imaging [24] and image deconvolution [25], in Sec. 2 we explore in greater detail the nature of spatial resolution determination using power spectral densities from X-ray fluorescence images. We show there that the spatial resolution at a given signal-to-noise ratio (SNR) depends on how one processes the signal and background.

### 1.3 Absolute photon flux in the two scanning stations

One aspect of characterizing the two SFXM stations was to measure excitation-dependent absolute photon flux Φ (*E*). The APS ran in top-up mode with a constant 100 mA of electron beam current in the storage ring, and we adjusted the beamline optics to maximize the flux observed via an ion chamber just downstream of the beam-defining pinhole at 48.15 m from the source. We carried out absolute photon flux measurements at 10 keV photon energy using a calibrated silicon PIN photodiode (Hamamatsu S3590-06) immediately downstream of each optic’s focus. We fed these signals into a low-noise current preamplifier (Stanford Instruments SR570); we used a voltmeter to confirm the measurement scale while the signal was passed to a voltage-to-frequency converter (NOVA N101VTF) for time-dependent measurements using 8-BM-B’s data acquisition system. To get the most reliable readings, we tuned the gain sensitivity *g*_s_ of the preamplifier to 2 µA/ V for the KB mirror, and 1 µA / V for the capillary. For this particular photodiode, 1 pA through the device corresponded to about 2382 photons/s at (*E*) = 10 keV, giving a photon flux-to-photocurrent conversion factor of *K* (*E*) = 2382 photons/(pA · s). Assuming negligible dark current voltage, we determined the absolute incident photon flux via

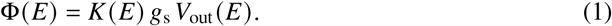

In this way, we found that the KB mirror station, at *E* = 10 keV, delivered a focused flux of Φ = 2.1× 10^10^ photons/s, while the capillary optic delivered a flux of Φ = 7.7×10^9^ photons/s (in each case, we estimated the absolute uncertainty to be approximately ±5% based on slow variations in the measurements).

### 1.4 Fluorescence data collection

While there are a variety of sample orientations and detector configurations that can be used in SFXM [4, 26, 27], most experiments mount the X-ray fluorescence detector at 90° to the incident beam direction so as to minimize elastic and inelastic scattering from horizontally polarized beams at synchrotron light sources [28]. In addition, the sample and scanning stage motions are typically inclined to the normal of the incident beam by 15° or 45° for a compromise between minimizing X-ray fluorescence self-absorption in planar samples while not introducing too much effective beam broadening in the scanning direction, as illustrated in Fig. 3. The effective beam width 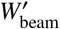 along the scanning stage direction, relative to beam width *W*_beam_ on a sample oriented at normal incidence to the beam, is given by

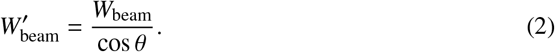

As noted above, we used *θ* = 15° and *θ* = 45° for the capillary and KB optics, respectively.

All data shown here were taken with continuous scanning, or “fly scan” mode, where the specimen was moved at near-constant velocity along the fast axis of the scan raster, with data recording advancing over time increments corresponding to position increments. With the KB scanning system, we frequently examined a large sample area using Δ_*x*_ = Δ_*y*_ = 25 µm, and then acquired higher resolution scans using Δ_*x*_ = Δ_*y*_ = 5-25 µm. With the capillary optic, we typically used Δ_*x*_ = Δ_*y*_ = 1-20 µm. In both cases, we chose per-pixel imaging times of *t*_dwell_ = 20-100 ms. With an incident X-ray beam energy of 10 keV, both fluorescence detectors were able to measure the spatially-resolved concentration of biologically-important elements including phosphorus, potassium, calcium, iron, nickel, copper, and zinc.

In order to determine the distance from the sample to the X-ray fluorescence detector and thus determine the solid angle of collection Ω, we collected fluorescence intensities *I* Δ*d* at three different detector displacements (Δ*d*) of 0, 3, and 6 mm. Because the solid angle, therefore the detected intensity, varies according to an inverse square law, we were able to determine the nominal sample-to-detector distance *d*_sd_ using

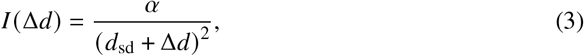

where *α* is a fitting constant. We used an ion chamber to correct for relative variations in incident flux, and we used the strong calcium fluorescence line from the sample shown in Fig. 1 as the signal. This gave us detector distances of *d*_sd_ = 31.2 mm for the KB system and *d*_sd_ = 14.1 mm in the capillary system. Only three of the four detector elements in the KB system were functioning, reducing its active area from *A*_active_ = 170 mm^2^ to 127.5 mm^2^. With the 7-element detector in the capillary system, the active area was reduced from *A*_active_ = 350 mm^2^ to 280 mm^2^ by the use of a collimator that minimized the sensitivity to scattering and stray fluorescence from materials other than the sample area in the beam focus. From these values, we were able to determine that the solid angles of collection Ω, which we calculated via

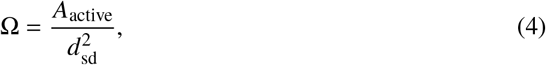

were 0.13 sr and 1.46 sr for the KB and capillary systems, respectively.

The Xspress3X detector readout electronics used with the 7-element detector for the capillary optic scanning station can store one scan line work of data internally, and this required required modifications to the data acquisition software. To increase data collection efficiency, we developed a graphical user interface (GUI). This software allows users to define multiple scans that can be executed sequentially as part of integration the Xspress3X readout system with the APS’ EPICS controls system. The batch scanning software prompts the user for scan parameters such as scan type, scan width, number of points per line, dwell time, and the center position. It verifies that the parameters for each scan do not exceed motor position or velocity limits, and returns an estimated completion time if the parameters are valid. At the start of each scan in the queue, the software resets the acquisition hardware and readies the controls hardware such that every *n*th motor pulse produces a single trigger event based on the ratio of the desired pixel size to motor resolution. This ensures that each trigger event captured on the Xspress3X is equidistant from the last, resulting in uniform pixel size. This new controls software differs from the one used for the XIA XMAP readout system used with the 4-element detector for the KB mirror scanning station in that it includes features such as parameter import/export functionality, event logging, and parameter validation. We also calibrated the Xspress3X system to ensure accurate and consistent measurement across each of the detector elements and electronics channels prior to commissioning.

### 1.5 Fluorescence data processing

Energy-dispersive X-ray detectors collect all the charge from above-threshold photon detection events and use the 3.65 eV energy for electron–hole separation in silicon [29–31] to estimate the energy of fluorescent photons. However, the collected signal also includes photons that are elastically and inelastically scattered from the sample and also from instrument materials, incomplete charge detection, and escape peaks in the detector material [8]. Therefore, a variety of analysis programs have been developed to use the as-recorded signal and separate X-ray fluorescence photon events from the background [9–14]. In our case, we used two programs: MAPS [10] and M-BLANK [14]. MAPS offers both region-of-interest (ROI) selection (sometimes called “spectral binning”), and full-spectrum fitting algorithms. M-BLANK offers full-spectrum fitting. ROI selection is still used at some synchrotron beamlines, so we show in Supplementary Material Sec. S2 the advantages of using full-spectrum fitting codes like those offered by MAPS and M-BLANK.

## 2. Resolution determination via power spectral density

One commonly-used method for evaluating the spatial resolution of images is to use 3D Fourier shell correlations (or 2D Fourier ring correlations) between two independent images of the same specimen as a function of spatial frequency *u* = 1/ (spatial period) [32, 33]. However, expediency in data collection as well as radiation dose minimization make it useful to evaluate spatial resolution from single images. With low-exposure images, this can be carried out using power spectral density (PSD) analysis. In two dimensions, we calculated the (PSD) *S u*_*x*_, *u*_*y*_ for each element using

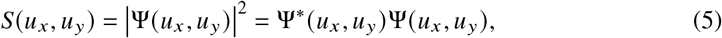

where

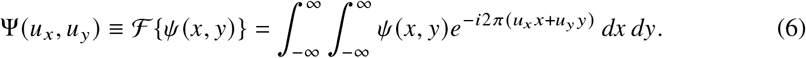

In the above, Ψ (*u*_*x*_, *u*_*y*_) is the Fourier transform of the real space X-ray wavefield *ψ* (*x, y*) leaving the object (the exit wave), which, in transmission imaging, can be found from the square root of the intensity if there is no appreciable phase contrast. In keeping with this convention for fluorescence as an incoherent process, we generated *ψ* (*x, y*) from the square root of each element’s fluorescence image. We note that with M-BLANK, to avoid negative square roots [14], we offset each pixel value by the largest negative value in the fluorescence image (constant offsets in an image affect only the zeroth spatial frequency in the Fourier transform). In the discrete Fourier transform, the spatial frequency *u*_*x*_ runs from −1/(2Δ_*x*_) to *b*_*x*_/(2Δ_*x*_) with *b*_*x*_ = (*N*_*x*_/2 − 1) /(*N*_*x*_/2), with a similar result for *u*_*y*_. In most cases, we show *S*(*u*_*r*_), where 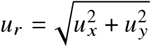. We can use a smaller number of steps in *u*_*r*_ than 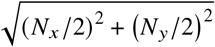 so as to smooth out the resulting power spectral density. To ensure that *u*_*r*_ = 0 was at the central pixel, and for efficient use of the fast Fourier transform (FFT), we padded images with odd numbers of rows or columns with average signal levels to lead to even numbers of rows and columns *N*_*x*_ and *N*_*y*_, respectively.

When shown on a logarithmic scale for power spectral density versus spatial frequency, *S*(*u*_*r*_) typically declines linearly with *u*_*r*_ for many types of images. This corresponds to a signal trend of

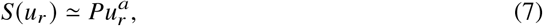

where *a* < 0. The magnitude of *a* has generally been thought to be somewhere between 3 and 4 [15]. As shown in Sec. 2.4, in some cases we obtain a larger slope (see Supplementary material Sec. S3 for an additional explanation on this). Next, when images are obtained using a finite number of photons with associated Poisson noise statistics, it is common for *S* (*u*_*r*_) to decrease until it reaches a constant “noise floor” value *S*_nf_. This is due to Poisson fluctuations being uncorrelated across pixels; as a result, Poisson noise is like a delta function in real space with a “flat” power spectral density that is uniform over all pixels in Fourier space. One can exploit these trends in signal and noise for Wiener noise suppression on Fourier plane representations of images, where the Wiener filter *W* (*u*_*r*_) is given by [34]

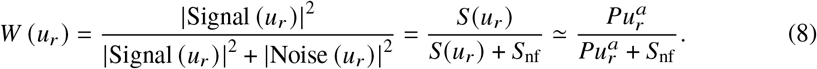

Because *W* (*u*_*r*_) ≃ 1 at low spatial frequencies where *S u*_*r*_ dominates, and because *W u*_*r*_ ≃ *0* at high spatial frequencies where the noise floor *S*_nf_ dominates, it is only in the “knee” region in power spectral density versus spatial frequency, where *S*(*u*_*r*_) ≳ *S*_nf_, that one requires an estimate of the trend of *S*(*u*_*r*_) for Wiener filtering. In this spirit, one can use a fit of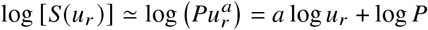 to find the power law decrease slope *a* for the signal.

That is, one can use data points fromS spatial frequencies *u*_*r*_ somewhat lower than the “knee” to find the signal trend and thus determine *P* and *a*. One can also use the average of data points at spatial frequencies *u*_*r*_ above the “knee” to find

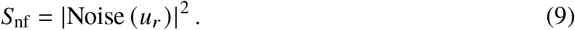

With *P, a*, and *S*_nf_ now determined, the spatial frequency *u*_knee_ of the “knee” is the value of *u*_*r*_ for which 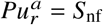, or

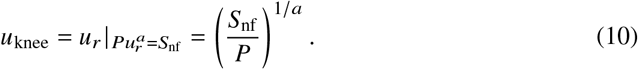

For evaluating the effective spatial resolution of an image, it is best not to have the signal be equal to the noise (the condition of the “knee”), but to have the signal be higher than the noise. It is common to use the Rose criterion [35] of requiring a signal-to-noise ratio of SNR_res_ = 5 for high-quality images (lower SNR values are often deemed acceptable in specific contexts). Thus, we can obtain an estimate of the maximum spatial frequency *u*_res_ corresponding to a high-quality image from the case where the resolution-defining power spectral density is a multiple SNR_res_ of the noise floor *S*_nf_, or

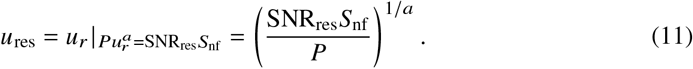

This leads to a spatial resolution estimate of

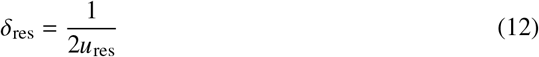

based only upon the power spectral density of a single image.

We developed a Python computer program (using a PyQt graphical user interface) called PyXRFPower to carry out power spectral density analysis on X-ray fluorescence images. The program allows one to either estimate the spatial resolution assuming a symmetric beam profile or the resolution in the horizontal and vertical directions (Sec. 2.3). After reading in a scanning fluorescence X-ray microscope dataset, one can select images corresponding to emission at one or several elemental X-ray fluorescence lines and carry out power spectral density fitting separately for each element. The selection of the data points to use for the signal trend *S* (*u*_*r*_) (Eq. 7) and noise floor *S*_nf_ (Eq. 9) is accomplished by dragging a cursor through spatial frequency regions on either side of the “knee” in the power spectral density. Spatial frequencies *u*_res_ of Eq. 11 resulting from a user-defined value of SNR_res_ and corresponding spatial resolutions are reported alongside data trend fit slopes *a* as provided by Eq. 7. PyXRFPower is currently available on Github: https://github.com/bwr0835/pyxrfpower.

Using the approaches described above, we characterized the scanning fluorescence X-ray microscopy performance of both the KB mirror and capillary optic experimental stations. For the KB mirror, we acquired images with scan step sizes of Δ_*x*_ = Δ_*y*_ = 5 µm and per-pixel acquisition times of *t*_dwell_ = 25 ms (97.7% live time) and 100 ms (99.6% live time). For the capillary optic, we rotated the original sample 90° clockwise to match the orientation of the fields of view shown in Figs. 1(a) and (b), but we calculated resolution using the original orientation. The scan step size for the capillary optic was Δ_*x*_ = 1 µm and Δ_*y*_ = 2 µm, and we used *t*_dwell_ = 20 ms (100.0% live time) and 50 ms (99.6% live time). Unless stated otherwise, the results shown below were obtained using the M-BLANK program [14] for fluorescence spectrum analysis, as described in Sec. 1.5.

### 2.1 Spatial resolution versus fluorescing element

As noted in Sec. 1, the spatial resolution actually achieved in an image depends on the optic but also on the signal-to-noise characteristics of the X-ray fluorescence image [24, 25]. We therefore evaluated SFXM images separately for each element using the methods described in Sec. 2 to find the power spectral density trends for the signal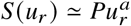 (Eq. 7) and the noise floor *S*_nf_ (Eq. 9). From that, we were able to obtain spatial resolutions *δ*_res_ (Eq. 12) for a signal-to-noise ratio of SNR_res_ = 5.

For the capillary optic, we imaged the specimen described in Sec. 1.1 with 339 × 242 pixels of size Δ_*x*_ = 1 µm and Δ_*y*_ = 2 µm with a per-pixel imaging time of *t*_dwell_ = 50 ms. Of all the elemental images obtained from this sample, the as-scanned calcium image shown in Fig. 4(a), which is the same capillary image shown in Fig. 1(c) but rotated, appeared to have the most favorable signal-to-noise ratio. Therefore we carried out power spectral density analyses on that Ca image plus images of Fe and Zn fluorescence. The results displayed in Fig. 4(b) show resolution results of *δ*_res_ = 6.3 µm for Ca, *δ*_res_ = 13.5 µm for Fe, and *δ*_res_ = 17.8 µm for Zn. The spatial resolution estimate for the Ca image was in good agreement with the spatial resolution estimated from images of a standard test pattern.

**Fig. 4.**
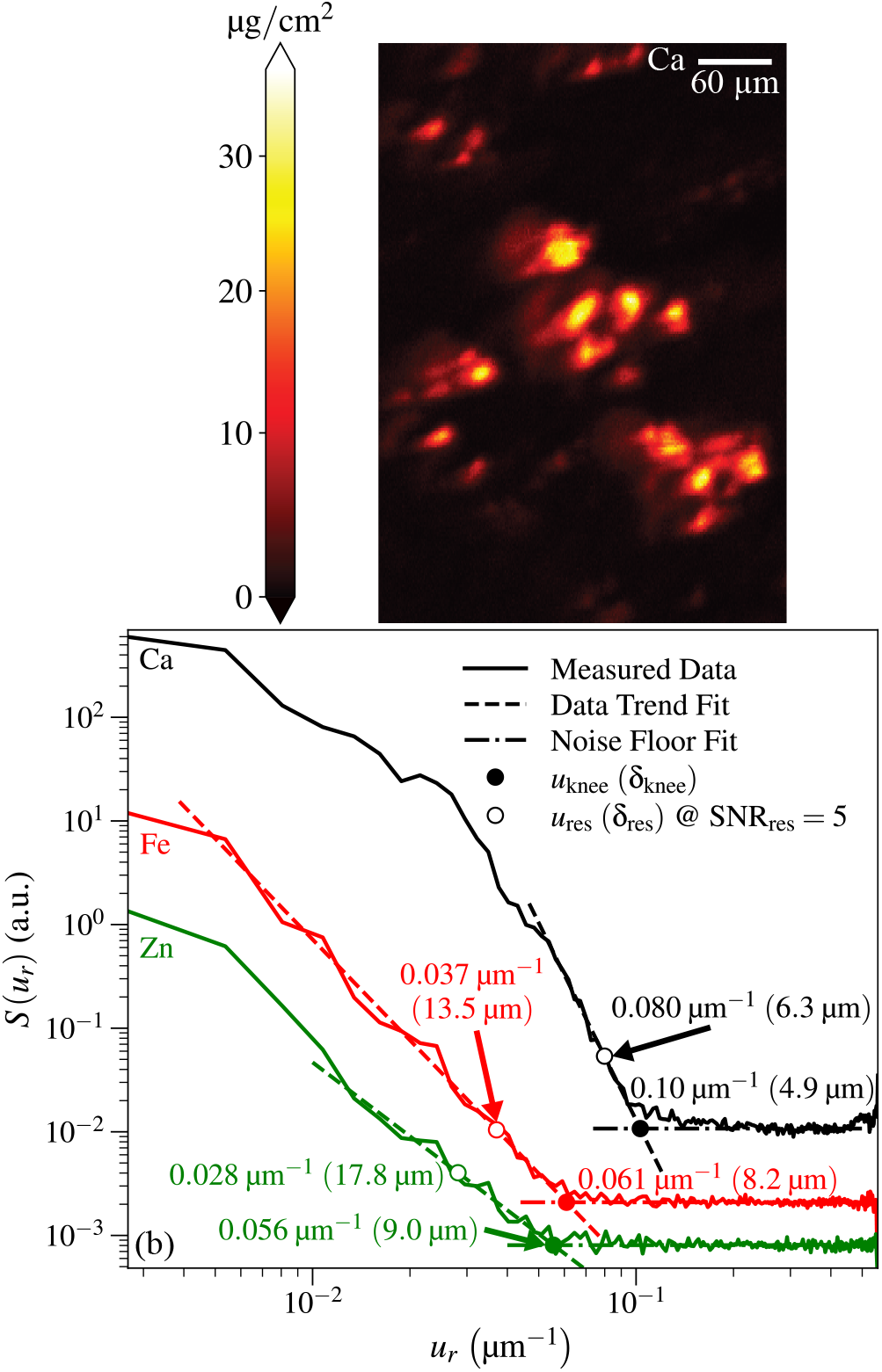
Image of a *Candida albicans*-infected mouse kidney partial section obtained using the capillary optic scan station and a per-pixel scan time of *t*_dwell_ = 50 ms. At top is the calcium fluorescence image from Fig. 1(c) as scanned, which is rotated relative to the orientation shown in that figure. At bottom (b) are power spectral density plots for the images of three different elements obtained from the same scan: Ca, Fe, and Zn. Because of differences in concentrations of each of these elements in the sample and differing fluorescence yields and background signals, each element’s image and power spectral density has different signal and noise characteristics. As described in Sec. 2.1, this leads to different values for spatial frequency *u*_knee_ (Eq. 10) and corresponding spatial resolution *δ*_knee_ of the “knee”, as well as the spatial frequency *u*_res_ (Eq. 11) and corresponding spatial resolution *δ*_res_ (Eq. 12) for a signal-to-noise ratio of SNR_res_ = 5.

We also scanned a larger area of a region in the same specimen, containing most of the capillary field of view, using the KB mirror scanning station. This scan was over 187 × 75 pixels of size Δ_*x*_ = Δ_*y*_ = 5 µm with a per-pixel imaging time of *t*_dwell_ = 100 ms. This pixel size was just small enough for meeting the Nyquist criterion, and Fig. S2 in Supplement 1 shows that it was small enough to allow the noise floor to be seen. Using the same analysis approach as shown in Fig. 4, we obtained resolution estimates of *δ*_res_ = 10.5 µm for Ca, *δ*_res_ = 12.7 µm for Fe, and *δ*_res_ = 12.9 µm for Zn.

The strong Ca fluorescence provided a good measure of the optic-limited resolution in both cases. The weaker Fe and Zn signals came closer to the optic limit in the longer-dwell-time KB station measurement than they did in the shorter-dwell-time capillary station measurement, further illustrating the point that the achieved spatial resolution depends both on the optic and on signal levels.

### 2.2 Achieved resolution versus scan time

As one increases the per-pixel exposure time *t*_dwell_, the signal *S* (*u*_*r*_) (Eq. 7) should increase relative to the noise floor *S*_nf_ (Eq. 9), meaning that spatial resolution *δ*_res_ (Eq. 12) should improve. For the Ca image obtained with the KB mirror optic, the resolution at *t*_dwell_ = 25 ms was *δ*_res_ = 14.7 µm, corresponding to a 58% worsening of spatial resolution compared to the result of *δ*_res_ = 10.5 µm at *t*_dwell_ = 100 ms. Because the spatial resolution limit involves both the intrinsic resolution of the optic and the signal-to-noise ratio, it does not scale simply with exposure time alone. For the Ca image obtained with the capillary optic, decreasing the per-pixel exposure time to *t*_dwell_ = 20 ms gave *δ*_res_ = 8.5 µm compared to *δ*_res_ = 6.3 µm at *t*_dwell_ = 50 ms, or about a 40% worsening of spatial resolution. These results clearly show how the achieved spatial resolution depends strongly on exposure time for photon statistics-limited images such as are obtained in scanning fluorescence X-ray microscopy of intrinsic concentrations of metals in biological tissues. See Fig. 5 for a figure of the two *S* (*u*_*r*_) profiles we obtained for the capillary at the two values of *t*_dwell_ we used.

**Fig. 5.**
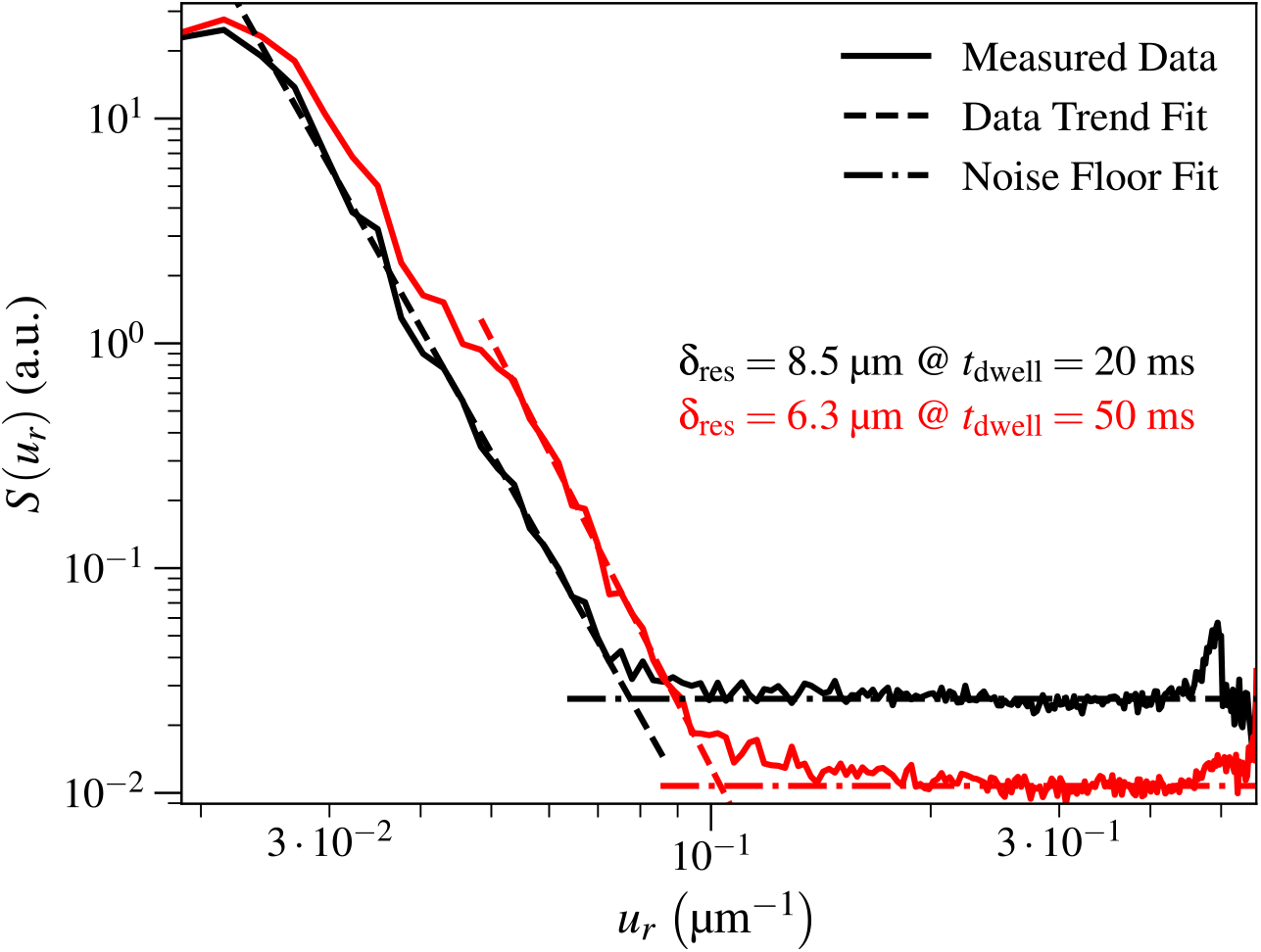
Comparison of isotropic capillary spatial resolutions *δ*_res_ when increasing per-pixel exposure time *t*_dwell_. The resolution improved from *δ*_res_ = 8.5 µm to *δ*_res_ = 6.3 µm when increasing *t*_dwell_ from 20 ms to 50 ms since the incident photon fluence ℱ, which affects signal and noise levels, increases in direct proportion with *t*_dwell_ (see Eq. 14).

### 2.3 Non-azimuthally-symmetric spatial resolution estimate

Up to this point, we have assumed that the spatial resolution is the same in the horizontal and vertical directions. However, it is not uncommon at synchrotron light source beamlines to have slight spatial resolution asymmetries due to both beamline optics and tilting the sample and scanning stage motion as shown in Fig. 3. One can therefore modify the above analysis so that it is carried out in two distinct azimuthal angle ranges as shown in Fig. 6, with azimuthal angle ranges of ± 30° about each direction representing one reasonable choice. This allows for resolution estimates of *δ*_res,*x*_ and *δ*_res,*y*_, respectively.

**Fig. 6.**
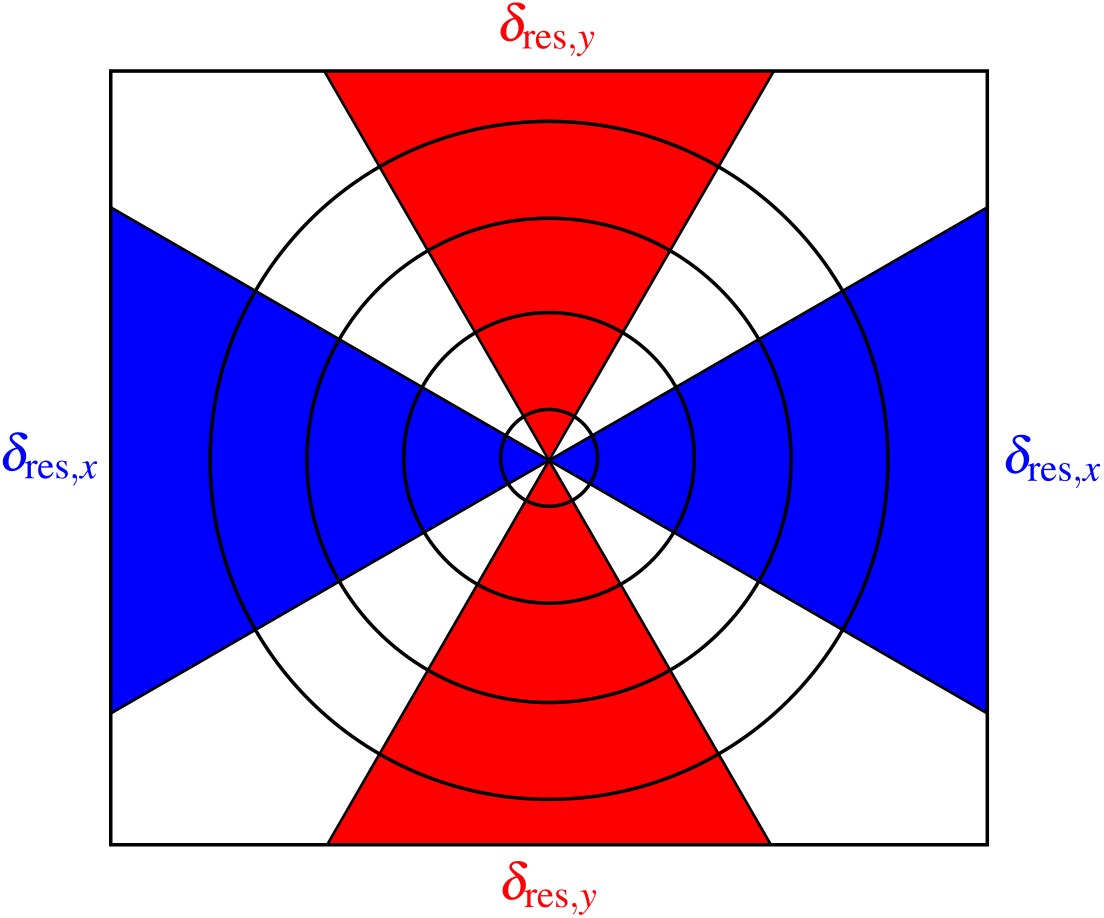
When images have asymmetry in their spatial resolution, one can define azimuthal angle ranges about the *x* and *y* axes (shown here as blue and red regions, respectively) to obtain separate estimates of spatial resolution of *δ*_res, *x*_ and *δ*_res,*y*_. We used azimuthal angle ranges of ±30° about the *x* and *y* axes.

For the Ca image shown in Fig. 4(a) for which the azimuthally-averaged resolution was *δ*_res_ = 6.3 µm, we also used the approach shown in Fig. 6 to obtain separate estimates for the spatial resolution in the *x* and *y* directions, yielding *δ*_res,*x*_ = 6.6 µm and *δ*_res,*y*_ = 6.1 µm as shown in Fig. 7. We note that because the maximum value of *u*_*x*_ was decreased by a factor of two due to using Δ_*x*_ = 2 µm step size (versus Δ_*y*_ = 1 µm), *S*_*x*_ (*u*_*r*_) does not extend out to the same value of *u*_*r*_ as *S*_*y*_ (*u*_*r*_). For the KB mirror, where the azimuthally-averaged Ca image resolution was *δ*_res_ = 10.5 µm, the resolution in the two directions was *δ*_res,*x*_ = 9.4 µm and *δ*_res,*y*_ = 12.2 µm.

**Fig. 7.**
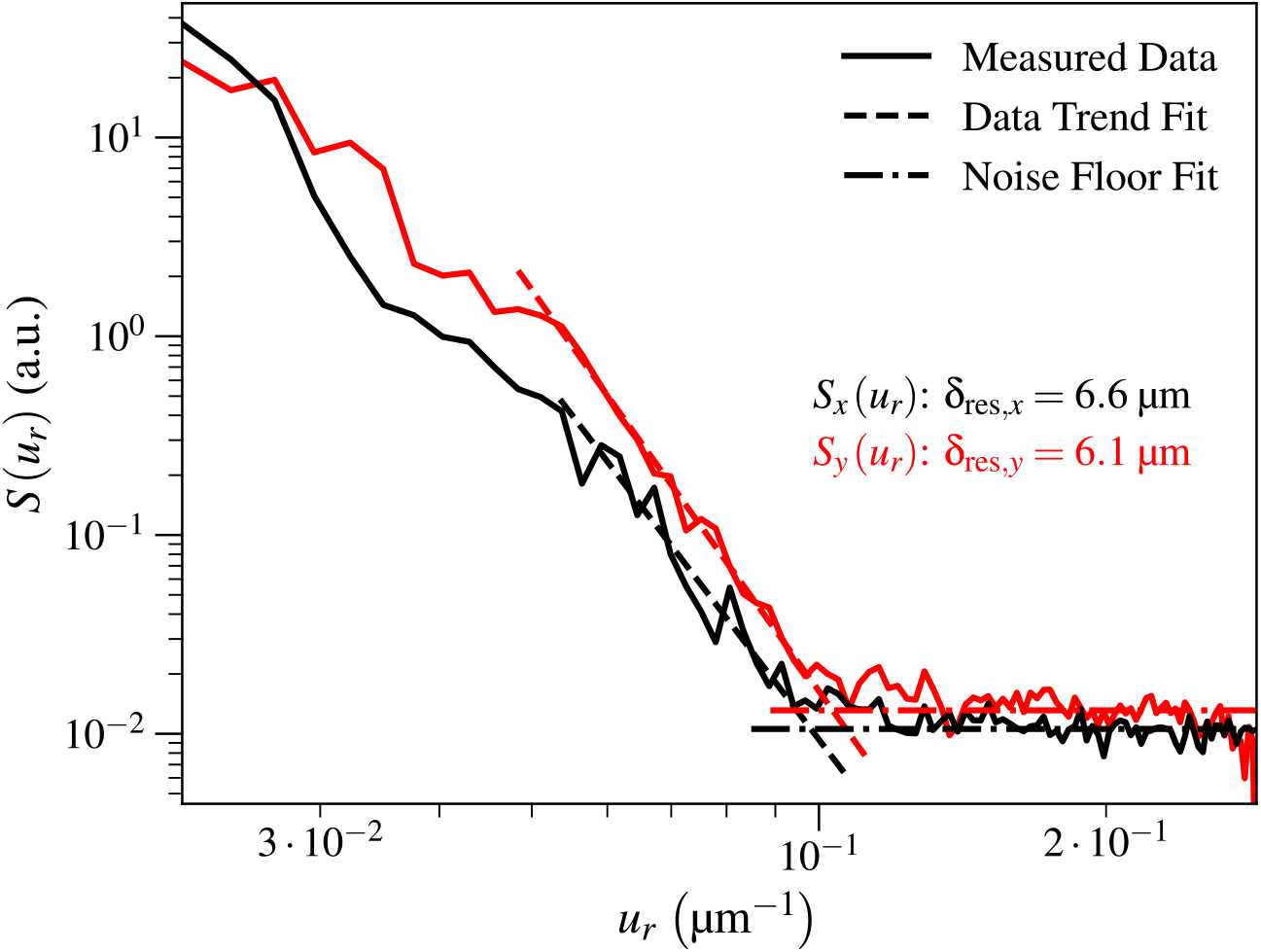
Using the same Ca image as shown in Fig. 4(a), we separately calculated power spectral densities *S*_*x*_ (*u*_*r*_)and *S*_*y*_ (*u*_*r*_) with the method shown in Fig. 7 with angular ranges of ±30° about the *x* and *y* axes, respectively. This yielded spatial resolution estimates of *δ*_res, *x*_ = 6.6 µm and *δ*_res,*y*_ = 6.1 µm, whereas the azimuthally-averaged spatial resolution result in Fig. 4 was *δ*_res_ = 6.3 µm. We truncated the maximum value of *u*_*r*_ shown here to 0.29 µm^−1^. One possibility for the spatial resolution mismatch is that there was a slight tilt misalignment of the capillary optic.

### 2.4 Sensitivity in resolution due to point selection

Our method of evaluating spatial resolution using power spectral density in photon-limited images relies on fits of the “signal” region of the power spectral density using 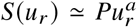 (Eq. 7), and the “noise” floor *S*_nf_ (Eq. 9). As noted above, the “signal” points should be selected over some range of spatial frequencies below the “knee” in the power spectral density profile at *u*_knee_ (Eq. 10), while the “noise” floor should be obtained by averaging points above *u*_knee_. In order to test the sensitivity of the spatial frequency *u*_res_ (Eq. 11) and the resulting resolution estimate *δ*_res_ of Eq. 12 to user selections, we show the results for three different selections of “signal” and the resulting values of slope *a* and resolution *δ*_res_ in Fig. 8. For the Ca image of Fig. 4(a), we obtained rather different slopes of *a* = −5.34, *a* = −6.32, and *a* = −6.73 with respective resolution estimates of *δ*_res_ = 6.19 µm, *δ*_res_ = 6.26 µm, and *δ*_res_ = 6.36 µm. While these slopes were a bit deviated from each other, the spatial resolution estimates spanned a 2.71% range. For the KB mirror field of view we acquired for *t*_dwell_ = 100 ms, we got *δ*_res_ = 10.52 µm, *δ*_res_ = 10.48 µm, and *δ*_res_ = 10.23 µm, and slopes of corresponded to *a* = −5.11, *a* = −5.57 and *a* = −6.59, respectively; these spatial resolution estimates spanned a range of 2.84%. We therefore infer that this approach allows one to estimate the achieved spatial resolution in a X-ray fluorescence image with a reproducibility of better than 3%. One might see greater variations in estimated resolution *δ*_res_ depending on “signal” trend points selected if there are both weaker signal and noise stabilities and weaker fluorescence signal and noise levels overall.

**Fig. 8.**
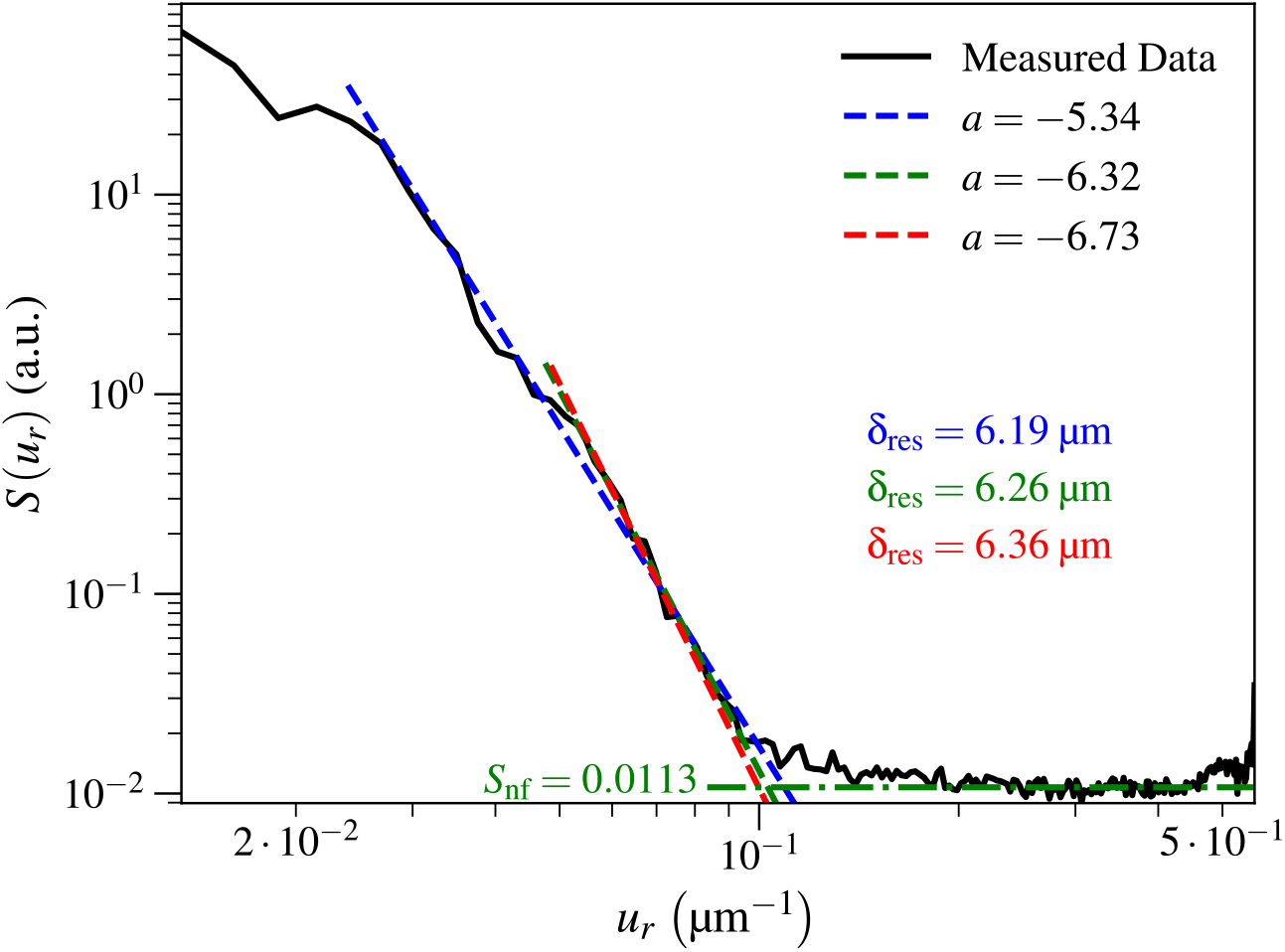
Test of the dependence on the estimated spatial resolution *δ*_res_ based on three different selections of the “signal” trend points to include in the fit of 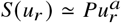 (Eq. 7), as discussed in Sec. 2.4. We carried this test out on the Ca image shown in Fig. 4(a). These three different estimates for the spatial resolution *δ*_res_ (Eq. 12) varied over a range of 2.71%.

## 3. Fluence per time on samples

The achievable spatial resolution in low-photon statistics images is limited in part by the fluence ℱ on the sample, or the cumulative photons per area [15, 24, 25], as shown in Fig. 5. We can assume that the area of the probe *A*_beam_ is given by

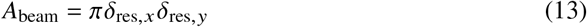

since, for an Airy probe, the spatial resolution is equal to the radii associated with the first minimum. With that in mind, we can use the absolute photon fluxes measured in Sec. 1.3 and the best resolution as obtained from Ca images (Sec. 2.3) to estimate the fluence per time *d*ℱ(*E*)/*dt a*s

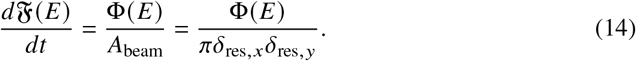

Using this approach, we obtained fluence per time estimates of *d*ℱ / *dt* = 5.8× 10^7^ photons / (µm^2^ ·s) ±5.8% for the KB mirror optic and *d*ℱ/ *dt* = 6.1 ×10^7^ photons / (µm^2^·s) ± 5.8% for the capillary.

## 4. Conclusion

We have modified the 8-BM-B beamline at the Advanced Photon Source (APS) at Argonne National Laboratory to provide two different setups for scanning fluorescence X-ray microscopy (SFXM) studies of intrinsic metals in biological tissue sections. The KB mirror station is best used for larger area scans at lower spatial resolution, while the prototype capillary optic station provides roughly a twofold improvement in spatial resolution. We have described the use of power spectral density analysis of single images to obtain spatial resolution estimates, and we have shown how the achieved spatial resolution is affected by signal strength from different fluorescing elements.

The results we reported in this work were for the beamline as it existed right before the shutdown of the original APS storage ring in April 2023. The upgrade of the APS (APS-U) [19] should offer a slight improvement in the brightness of bending magnet sources (and 75-100× improvement for undulator sources). Further gains might be possible by addressing limitations in the beamline toroidal mirror discussed in Sec. 1.2. Therefore, this beamline should be an even more valuable resource for future studies of elemental distribution in biological tissues, aiding user communities including that of the NIH-supported QE-MAP center at Michigan State University.

## Acknowledgments

The authors thank Professor Brendan Cormack (School of Medicine, Johns Hopkins University) for contributions in mouse infection studies, and Dr. Niharika Sinha (College of Natural Science, Michigan State University) for providing the visible light image of the infected mouse kidney used for the simulation discussed in Supplement 1.

## Data Availability Statement

Data underlying the results presented in this paper are currently available upon request and will be available on Dryad upon receiving a publication number.

## Supplemental Document

See Supplement 1 for supporting content.

## References

1. R. McRae, P. Bagchi, S. Sumalekshmy, and C. J. Fahrni, “In situ imaging of metals in cells and tissues,” Chem. Rev. 109, 4780–4827 (2009).

2. D. Z. Zee, K. W. MacRenaris, and T. V. O’Halloran, “Quantitative imaging approaches to understanding biological processing of metal ions,” Curr. Opin. Chem. Biol. 69, 102152 (2022).

3. J. Kirz, “Specimen damage considerations in biological microprobe analysis,” in Scanning Electron Microscopy, vol. 2 (SEM Inc., Chicago, 1980), pp. 239–249.

4. C. J. Sparks, Jr., “X-ray fluorescence microprobe for chemical analysis,” in Synchrotron Radiation Research, H. Winick and S. Doniach, eds. (Plenum Press, New York, 1980), chap. 14, pp. 459–512.

5. M. J. Pushie, N. J. Sylvain, H. Hou, et al., “X-ray fluorescence microscopy methods for biological tissues,” Metallomics 14, mfac032 (2022).

6. P. Horowitz and J. A. Howell, “A scanning x-ray microscope using synchrotron radiation,” Science 178, 608–611 (1972).

7. K. W. Jones, B. M. Gordon, A. L. Hanson, et al., “Application of synchrotron radiation to elemental analysis,” Nucl. Instruments Methods Phys. Res. B 3, 225–231 (1984).

8. R. A. Van Grieken and A. A. Markowicz, Handbook of X-ray Spectrometry, vol. 29 of Practical Spectroscopy (Marcel Dekker, New York, 2002), 2nd ed.

9. C. G. Ryan, “Quantitative trace element imaging using PIXE and the nuclear microprobe,” Int. J. Imaging Syst. Technol. 11, 219–230 (2000).

10. S. Vogt, “MAPS: a set of software tools for analysis and visualization of 3D x-ray fluorescence data sets,” J. de Physique IV 104, 635–638 (2003).

11. V.A. Solé, E. Papillon, M. Cotte, et al., “A multiplatform code for the analysis of energy-dispersive x-ray fluorescence spectra,” Spectrochimica Acta B 62, 63–68 (2007).

12. T. Schoonjans, L. Vincze, V.A. Solé, et al., “A general Monte Carlo simulation of energy dispersive x-ray fluorescence spectrometers – Part 5. Polarized radiation, stratified samples, cascade effects, M-lines,” Spectrochimica Acta B 70, 10–23 (2012).

13. T. Schoonjans, V.A. Solé, L. Vincze, et al., “A general Monte Carlo simulation of energy-dispersive x-ray fluorescence spectrometers – Part 6. Quantification through iterative simulations,” Spectrochimica Acta B 82, 36–41 (2013).

14. A. M. Crawford, A. Deb, and J. E. Penner-Hahn, “M-BLANK: a program for the fitting of x-ray fluorescence spectra,” J. Synchrotron Radiat. 26, 497–503 (2019).

15. C. Jacobsen, X-ray Microscopy (Cambridge University Press, Cambridge, UK, 2020).

16. M. Eriksson, J. F. van der Veen, and C. Quitmann, “Diffraction-limited storage rings – a window to the science of tomorrow,” J. Synchrotron Radiat. 21, 837–842 (2014).

17. P. Kirkpatrick and A. V. Baez, “Formation of optical images by x-rays,” J. Opt. Soc. Am. 38, 766–774 (1948).

18. A. S. Wildeman, N. K. Patel, B. P. Cormack, and V. C. Culotta, “The role of manganese in morphogenesis and pathogenesis of the opportunistic fungal pathogen candida albicans,” PLOS Pathog. 19, 1–26 (2023).

19. J. Kerby, “The Advanced Photon Source Upgrade: A brighter future for x-ray science,” Synchrotron Radiat. News 36, 26–27 (2023).

20. G. Kirker, S. Zelinka, S.-C. Gleber, et al., “Synchrotron-based x-ray fluorescence microscopy enables multiscale spatial visualization of ions involved in fungal lignocellulose deconstruction,” Sci. Reports 7, 41798 (2017).

21. S. L. Zelinka, J. E. Jakes, J. Tang, et al., “Fungal–copper interactions in wood examined with large field of view synchrotron-based x-ray fluorescence microscopy,” Wood Material Sci. Eng. 3, 174–184 (2018).

22. L. Copeland-Hardin, T. Paunesku, J. S. Murley, et al., “Proof of principle study: synchrotron x-ray fluorescence microscopy for identification of previously radioactive microparticles and elemental mapping of FFPE tissues,” Sci. Reports 13, 7806–7815 (2023).

23. M. Broda, J. E. Jakes, L. Li, et al., “Conservation of model degraded pine wood with selected organosilicons studied by XFM and nanoindentation,” Wood Sci. Technol. 58, 649–675 (2024).

24. J. Deng, D. J. Vine, S. Chen, et al., “Opportunities and limitations for combined fly-scan ptychography and fluorescence microscopy,” Proc. SPIE 9592, 95920U (2015).

25. J. Deng, D. J. Vine, S. Chen, et al., “X-ray ptychographic and fluorescence microscopy of frozen-hydrated cells using continuous scanning,” Sci. Reports 7, 445 (2017).

26. C. G. Ryan, R. Kirkham, R. M. Hough, et al., “Elemental x-ray imaging using the Maia detector array: the benefits and challenges of large solid-angle,” Nucl. Instruments Methods Phys. Res. A 619, 37–43 (2010).

27. Y. Sun, S.-C. Gleber, C. Jacobsen, et al., “Optimizing detector geometry for trace element mapping by x-ray fluorescence,” Ultramicroscopy 152, 44–56 (2015).

28. T. G. Dzubay, B. V. Jarrett, and J. M. Jaklevic, “Background reduction in x-ray fluorescence spectra using polarization,” Nucl. Instruments Methods 115, 297–299 (1974).

29. W. Shockley, “Problems related to p–n junctions in silicon,” Solid-State Electron. 2, 35–67 (1961).

30. B. G. Lowe and R. A. Sareen, “A measurement of the electron–hole pair creation energy and the Fano factor in silicon for 5.9 keV x-rays and their temperature dependence in the range 80–270 K,” Nucl. Instruments Methods Phys. Res. A 576, 367–370 (2007).

31. M. N. Mazziotta, “Electron–hole pair creation energy and Fano factor temperature dependence in silicon,” Nucl. Instruments Methods Phys. Res. A 584, 436–439 (2008).

32. W. O. Saxton and W. Baumeister, “The correlation averaging of a regularly arranged bacterial cell envelope protein,” J. Microsc. 127, 127–138 (1982).

33. M. v. Heel and M. Schatz, “Fourier shell correlation threshold criteria,” J. Struct. Biol. 151, 250–262 (2005).

34. W. H. Press, S. A. Teukolsky, W. T. Vetterling, and B. P. Flannery, Numerical Recipes: The Art of Scientific Computing (Cambridge University Press, Cambridge, UK, 1986).

35. A. Rose, “Unified approach to performance of photographic film, television pickup tubes, and human eye,” J. Soc. Motion Pict. Eng. 47, 273–294 (1946).

