## Supplementary Material for "Multifunctional bending magnet beamline with a capillary optic for X-ray fluorescence studies of metals in tissue sections"

### 1. Characterization of capillary focusing capability

In Sec. 1.2 of the main manuscript, we noted that the capillary optic's resolution with optimum illumination was much better than what was observed using the illumination available at APS (Advanced Photon Source) beamline 8-BM-B as it existed before the APS-Upgrade (the Upgrade is discussed in [1]). We outline here the optical tests of the capillary carried out using the small source size and divergence of undulator beamline 28-ID-B at the APS. The schematic of the experimental setup is shown in Fig. S1(a), and the capillary parameters are listed in Table S1.

APS beamline 28-ID-B uses an undulator with a period of 3.3 cm and a total length of 2.3 m. We employed a double-multilayer monochromator (DMM) and tuned it to an energy of 13 keV with a spectral bandwidth of 0.01%. The white beam slit, with dimensions of (0.1 mm)×(0.1 mm), functioned as the source. We used a polymer compound refractive lens (CRL) to image the white beam slit to a focal plane at 0.1 m downstream. According to prior quality measurements of the polymer CRL [2], the focal spot (acting as the secondary source) was estimated to be smaller than 1 μm when considering the geometric demagnification and the CRL aberrations. We positioned the test capillary 2.0 m downstream from the secondary source, with the expected focal plane located about 45 mm beyond the capillary center. This configuration aimed to produce a geometrical image size under 25 nm, predicated on a secondary source dimension of 1 μm and the assumption of ideal focusing optics. Consequently, any focal size measurements surpassing this threshold would likely result from aberrations within the capillary. We captured beam profiles using a detector system composed of a 100 μm thick LuAG:Ce scintillator, a 10× objective lens, and a Zyla sCMOS camera (Andor); this scintilator–camera system achievs an effective pixel size of 0.65 μm and a spatial resolution of 2.2 μm when accounting for the detector point spread function.

We identified the optimal focal spot to be at $z = 43$ mm downstream from the capillary center, with a FWHM size of (3.6 μm)×(3.7 μm), as shown in Fig. S1(b). Assuming that this beam size was combined in a RMS sense with the spatial resolution of the scintillator–camera system, we estimate the actual focal spot size of the capillary optic to be (2.9 μm)×(3.0 μm) FWHM, assuming a Gaussian beam distribution. This estimated size effectively represents the point spread function of the capillary. The primary factor contributing to the focal spot size was the capillary's internal surface slope error, with an average RMS slope error of 14 μrad. These results were consistent with predictions made using the surface figure data provided by the capillary manufacturer. In Figs. S1(c) and S1(d), we show the beam profiles captured at positions $z = 44$ mm and $z = 94$ mm downstream from the capillary center, respectively. The observed three-fold symmetry arises from the capillary's support structure, which partially obstructs the beam. The circular striations visible in Fig. S1(d) provide clear evidence of figure errors on the internal surface of the

Table S1. Designed and factory-achieved capillary parameters.

| | Specification | Achieved |
|---|---|---|
| Source-to-focus distance (mm) | 2027 | 2027 |
| Semi-minor axis (mm) | 1.075 | 1.072 |
| Capillary length (mm) | 50 | 50.3 |
| Entrance aperture diameter (mm) | 0.785 | 0.786 |
| Exit aperture diameter (mm) | 0.425 | 0.424 |
| Working distance (mm) | 20.0 | 20.1 |

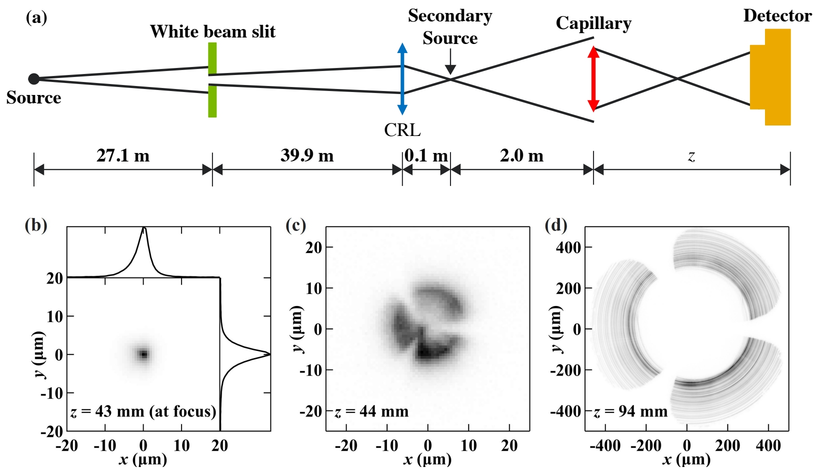


Fig. S1. Schematic of the 28-ID-B experimental setup (a) for measuring the focus capability of the capillary. Beam profiles were recorded at (b) $z = 43$ mm, (c) $z = 44$ mm, and (d) $z = 94$ mm from the downstream end of the capillary.

capillary.

### 2. Achieved resolution versus spectral analysis method

As noted in Sec. 1.5, the signal recorded by energy-dispersive detectors used in scanning fluorescence X-ray microscopy (SFXM) involves factors such as X-ray scattering directly into the detector, scattering onto instrument materials which in turn emit fluorescence photons, and incomplete collection of the charge from a photon absorbed in the detector. The most basic approach to fluorescence signal analysis is simply to define spectral regions of interest around each fluorescence peak, and to then treat the number of fluorescence photons detected as the total photon count within this spectral ROI (for example, a region covering the FWHM of a peak). This approach makes the simplifying assumption that the background is negligible compared to the photon count from strong fluorescence lines. However, most scanning fluorescence X-ray microscopy (SFXM) experiments now make use of spectral fitting methods as described in Sec. 1.5, where both signal and background distributions are estimated, so that the background can be subtracted from the signal.

The analyzed elemental signal values reflected in both $S(u_r)$ (Eq. 5) and its power law fit $Pu_r^a$ (Eq. 7), as well as the estimated noise floor $S_{\rm nf}$ (Eq. 9), can be affected by the analysis method used. When using signal trend and noise floor estimates to obtain a measure of the spatial resolution $\delta_{\rm res}$ (Eq. 12), this can affect the estimate of spatial resolution. In Fig. S2, we show the power spectral density for the same Ca dataset measured with a short per-pixel acquisition time of $t_{\rm dwell} = 25$ ms (thus, enhancing the challenges of background estimation) using the KB scanning station, but analyzed using two different approaches: the spectral ROI option as is available in MAPS [3] versus full-spectrum fitting as offered by programs like MAPS and M-BLANK [4] (we used M-BLANK in this case). This figure shows that the full-spectrum fitting approach led to a resolution estimate of $\delta_{\rm res} = 14.7$ µm, while the spectral ROI approach lead to a resolution estimate of $\delta_{\rm res} = 24.5$ µm. The differences between the achieved resolution values were quite significant, highlighting the advantages provided by full spectrum fitting.

### 3. Deviations in scanning probe profile

In $S(u_r) \simeq Pu_r^a$ (Eq. 7), as mentioned in Sec. 2 of the main manuscript, the value of the data trend slope magnitude $|a|$ in prior literature has generally been thought to be somewhere between 3 and 4 [5]; however, we obtained $|a| > 4$ for several different $S(u_r)$ profiles, including for the three profiles shown in Fig. 8 of that same document. Many experiments and theoretical models assume a focal spot of high contrast, meaning that there is very little flux in the regions outside the central focus (outside the Airy disk for a perfect circular optic). This applies to the case of most visible light optics, as well as Fresnel zone plates used as X-ray focusing optics. However, with grazing incidence X-ray optics, one often has a "halo" around the central focus due to imperfections in the figure and finish of the optics.

To see what effects these imperfections might have on $S(u_r)$, we took an area of a visible light microscope image of a healthy mouse kidney section (prepared by Niharika Sinha in the O'Halloran research group at

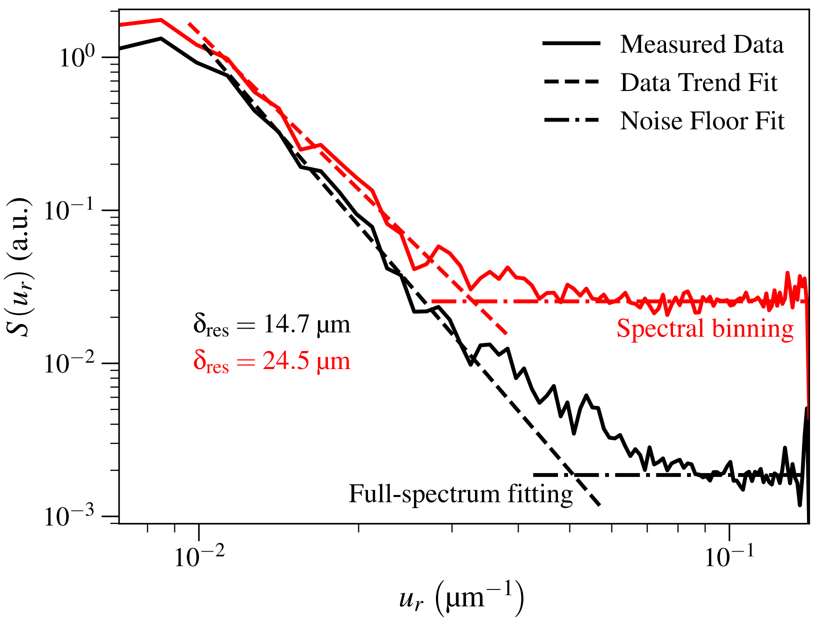


Fig. S2. Effect of different approaches to X-ray fluorescence spectrum analysis on the achieved spatial resolution as evaluated using power spectral density. Shown here are power spectral densities and fits for $S(u_r)$ (Eq. 7) and noise floor $S_{nf}$ (Eq. 9) for the Ca fluorescence dataset obtained using the KB mirror optic with $t_{dwell} = 25$ ms) with two different spectrum fitting approaches: spectral region-of-interest (ROI), or spectral binning (the signal is assumed to be the integral of what is measured over, for example, the FWHM peak width about each fluorescence emission line), and full-spectrum fitting applied to the spectrum observed by the energy-dispersive detector [4]. The estimate for achieved spatial resolution $\delta_{res}$ (Eq. 12) improved from $\delta_{res} = 24.5$ µm with the spectral binning method to $\delta_{res} = 14.7$ µm with the full-spectrum fitting method.

Michigan State University) and convolved it with probes representing two different point spread functions (PSFs). We used a visible light image to start due to the very significant photon counting statistics in it, so that it allowed us to simulate the effects of injecting Poisson noise later on. The first probe constituted a “sharp" spot of about one pixel size wide, as represented by a 2D delta function. The second probe focus constituted an X-ray mirror-type of PSF, where we added a Gaussian “halo" about ten pixels wide to an ideal single-pixel spot [5]. The resulting aggregate PSF ($\mathrm{PSF}_{total}$) was

$$\mathrm{PSF}_{total} = \delta\left(x,y\right) + \frac{A}{\sqrt{2\pi\sigma^2}} e^{-\left(x^2+y^2\right)/2\sigma^2}, \tag{S1}$$

where $\sigma$ is the standard deviation (or width) of the Gaussian, and where $A$ is a constant that was set to different fractions of the single-pixel probe energy $E_{sp} = 1$ to control the Gaussian strength. We tested $A = 0.01$, $A = 0.1$, and $A = 1$. After performing convolutions on the visible light image using each of these probes, we simulated the finite photon nature of an XRF elemental map by scaling each pixel intensity by the resulting image intensities and multiplying each of those fractions by a low number of photons. We then used a Poisson noise distribution to obtain “noisy” estimates based on the original signal level, thus simulating the Poisson noise usually present in SFXM images. We then calculated the resulting $S(u_r)$ profiles and their corresponding values for data trend fit slope $a$ and spatial resolution $\delta_{res}$. We normalized Eq. S1 by scaling $\mathrm{PSF}_{total}$ by its total energy $E_{total}$ for $A = 1$.

In Fig. S3, we show the resulting $S(u_r)$ profiles for a version of the ideal image with 100 photons per pixel in the brightest areas. In the cases where the “halo" contribution was low, we did not see a significant difference from the case of no “halo” around the sharp focus: the slopes were $a = -4.90$ for $A = 0.01$, and $a = -4.30$ for $A = 0.1$. However, when $A = 1$, we observed a power law slope of $a = -6.22$. We also obtained resolutions $\delta_{res} = 10.6$ µm and $\delta_{res} = 22.0$ µm for the $A = 0.1$ and $A = 1$ cases, respectively. From this, we can deduce that the profile of the probe used in an SFXM scan can lead to larger magnitudes of the slope $|a|$ in Eq. 7.

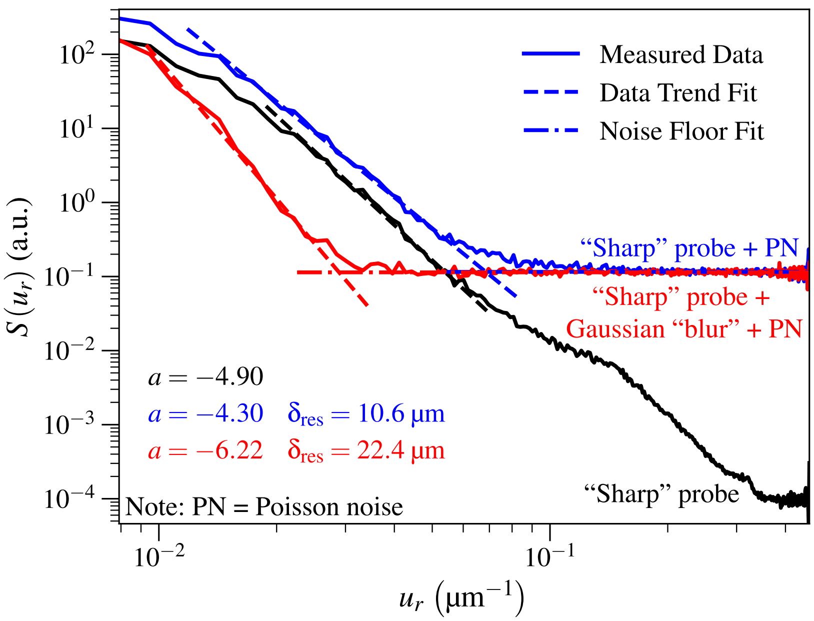


Fig. S3. Demonstration of how the scanning beam profile can affect the observed trend of signal versus spatial frequency of $S(u_r) \simeq P u_r^a$ (Eq. 7). We convolved a very low noise visible light image of a healthy mouse kidney section with an ideal, single-pixel illumination spot, as well as with a version of this spot (or probe) with an added Gaussian with $\sigma = 10$ pixels and a fraction of the energy $A = 0.01$, 0.1, and 1 of the single-pixel probe. This provides a way of approximating the "halo" effect one often has due to mirror surface imperfections with reflective X-ray optics. In the case of the largest probe halo fraction, the observed slope $a$ changed.

### References

1. J. Kerby, "The Advanced Photon Source Upgrade: A brighter future for x-ray science," Synchrotron Radiat. News **36**, 26–27 (2023).
2. Z. Qiao, X. Shi, P. Kenesei, *et al.*, "A large field-of-view high-resolution hard x-ray microscope using polymer optics," Rev. Sci. Instruments **91**, 113703 (2020).
3. S. Vogt, "MAPS: a set of software tools for analysis and visualization of 3D x-ray fluorescence data sets," J. de Physique IV **104**, 635–638 (2003).
4. A. M. Crawford, A. Deb, and J. E. Penner-Hahn, "M-BLANK: a program for the fitting of x-ray fluorescence spectra," J. Synchrotron Radiat. **26**, 497–503 (2019).
5. C. Jacobsen, *X-ray Microscopy* (Cambridge University Press, Cambridge, UK, 2020).
